# A broad sense measure of health and its properties in the HRS

**DOI:** 10.1101/081232

**Authors:** Phillip Cantu, Connor M. Sheehan, Benjamin W. Domingue

**Author notes:** Corresponding authors: Phillip Cantu, Department of Sociology and Population Research Center, University of Texas at Austin. Population Research Center, G1800, University of Texas at Austin, Austin, TX 78712-0544. Ben Domingue, Graduate School of Education, 508 CERAS Stanford CA 94304.

## Abstract

Measuring health is a crucial component of much social research. Two approaches are typical. Health may be measured via either narrowly targeted questions about aspects of disability or chronic conditions. Alternatively, individuals may be asked to self-report their health in some global sense. Both approaches have potential drawbacks. We consider a broad sense measure of health constructed by items from five different batteries related to physical and mental wellbeing. We demonstrate that this measure predicts time until death better than self-reported health, especially for females. Although this measure has promise, we argue that future surveys on health would benefit from the inclusion of additional items focusing on issues salient to younger individuals or other non-disabled respondents.

## 1. Introduction

Understanding health and its age-associated dynamics is a key objective of the Health and Retirement Study (HRS) and related studies (e.g., ELSA, MHAS, etc.). Crucial to this effort is a critical investigation of what researchers mean by “health” and how health is measured. Previous research has tended to either (a) look at specific, narrow domains of health (i.e. activities of daily living) or (b) ask respondents to self-report their overall health status. The first approach is common for the study of disability. For example, the battery of questions which comprise the Activities of Daily Living ask if a respondent has the ability to conduct basic daily tasks of functioning (1). While these measures are effective in measuring inability to perform tasks, they cover relatively restrictive domains and presumably do not provide information about relative differences between individuals who are not disabled and who may still have different amounts of “health.” There are likely substantial differences in overall health between respondents with the exact same reports or levels of ADLs, but this heterogeneity is not captured in research focusing on disability. The second approach, asking respondents to self-report their general or overall health, has been used in a vast amount of research (e.g., 2,3). Research has found that self-reported health corresponds with physician evaluations and accurately predicts mortality (4). This measure takes a more sweeping view of “health” than, say, the Activities of Daily Living battery, but this comes at the potential expense of introducing subjectivity into the measurement process. Individuals may respond differently to this question partially as a function of their age (5), sex, and ethnicity (6,7) which is undesirable.

While both of these approaches have been fruitful and will continue to be used to operationalize health, we ask whether a broader, yet still objective, consideration of what constitutes “health” can yield new insight. Our approach is atheoretical in the sense that we are endeavoring to make better use of valuable extant data. Our lack of a strict theoretical foundation for what is meant by “health” is similar to the manner in which self-reports of “health” are based on the respondents’ subjective ideas about what this might mean (e.g., does the respondent consider their mental health when they are asked generically about “health”). We comment on this purely utilitarian viewpoint in Section 4 after first demonstrating that it is offers potential utility. We use Item Response Theory (IRT; 8) to combine multiple batteries of questions related to functioning, chronic conditions, and psychological health to form a set of health-related items that we use to construct a broad sense measure of health on a continuous scale. We focus on two questions. *First*, what does the IRT analysis tell us about the ways in which our inquiries regarding health could be improved? *Second*, how does our broad sense measure of health potentially improve upon specific batteries and the standard self-reported health measure? Given the graying of the US population (9), these are timely questions.

### 1A. Current operational definitions of “health”?

In much demographic inquiry, researchers have attempted to measure health as a broad construct regarding overall wellbeing. Studies examining the relationship between educational attainment and “health” are frequently utilize such an indicator (10). Researchers use self-reported health SRH as a way to quickly measure global general health due to its simplicity, convenience, and also because it is has a well-documented history as a strong predictor of mortality and aligns well with doctor assessments of health (4,11,12). There are open questions about the validity of SRH, however. For example, there may be period effects related to how people think about health (13). Other research illustrates difficulty in making cross-ethnic comparisons, especially with relation to differences in linguistic backgrounds (6,7). The subjective nature of SRH is thus a cause for concern in certain types of inquiry (e.g., inquiry based on diverse populations).

Other types of research have focused on health as measured by different batteries of items inquiring about narrowly defined aspects of respondent wellbeing and functioning and have provided valuable information to researchers about the nature of physical decline (14–22). Broadly speaking, this research conceptualizes health as living without disability. The disablement process (1) offers a framing through which researchers can understand the multiple dimensions of health and wellbeing; specifically the disablement process shows that disability as measured by survey items are the downstream result of pathological conditions upstream. The disablement process allows researchers to conceptualize health in an objective manner (i.e. an individual responds that he or she is cannot walk across a room). Such a focus on disability is less appropriate for use in broader populations that contain younger respondents or healthier (non-disabled) older respondents.

### 1B. The Broad Sense Health approach

Given the potential subjectivity of SRH and the narrowness of other measures based on functional limitations and behavioral modifications, we think there is room for alternative operational definitions of health. Survey questions are a useful tool in health-related research, in part due to the fact that they are relatively cheap to administer (c.f., clinical assessments) and are included in many studies of human behavior. Survey questions on health can cover a broad variety of ground, from function (23) to diagnosed chronic conditions (24) and mental health (25). We combine information from across these survey items into a single IRT-based composite: broad sense health (BSH). Item response theory was originally developed for educational measurement, but these methods can also be used to study health and functioning. Previous work has utilized these techniques to study differences in response processes as a function of ethnicity on indicators of disability (26,27). IRT is a useful tool in this context in that it allows for both a holistic summary of health *and* a critical appraisal of measurement quality. We examine both issues in this paper. In a time with increasing non-response (28) and decreasing funds for surveys, knowing which questions can quickly differentiate the “healthy” from “unhealthy” is critically important.

## 2. Data & Methods

### 2A. Sample

We utilize the Health and Retirement Survey (24,29). The HRS is a nationally representative of the non-institutionalized US population longitudinal survey of older Americans. We use data from wave 4, collected in 1998, for our analysis because in 1998 two new cohorts were added which made the HRS nationally representative for those aged 50+ for the first time. The sample contained 21,384 respondents, all those respondents with at least some available information in 1998 on the health indicators. The respondents had a mean birth year of 1932 (SD=11.2; average age of 66 as of 1998). The majority of the sample (58.1%) was female. The sample was 76.3% non-Hispanic white, 13.9% non-Hispanic black, and 6.1% Hispanic. We compare our BSH measure to SRH which is measured in HRS as a 5-category scale (1-excellent, 5-poor). In our analytic sample 12.3% of respondents self-reported excellent health, 25.6% very good health, 30.6% good health, 20.6% fair health, and 10.9% poor health.

### 2B. Indicators

We use health indicators from the following batteries: Activities of Daily Living, Instrumental Activities of Daily Living, Functional Limitations, Depression, and Chronic Conditions. Additional information on the wording of individual items is contained in the Supplemental Information (SI). Items are coded so that “0” indicated that the respondent reported having a condition or limitation measured (“1” indicates better health in all cases).^1^ Information on the content of the utilized items can be found in the SI. Note that we focus on items related to both physical and mental health. We argue that SRH is also likely to reflect the influence of aspects of both physical and mental health. Here, we have the capacity to inquire about the relationship between different indicators (e.g., physical functioning and mental health) and the respondent’s overall health as measured by BSH.

Descriptive statistics for the indicators can be found in the first two columns of Table 1. For the average indicator, 83% of individuals did not have the health condition, physical limitation, depressive symptom, etc. of interest. Some items captured almost no variation in respondent health or behavior. Even the most prevalent of conditions/limitations was not prevalent in more than half of our sample (minimum prevalence of 0.52). This variation in prevalence is also shown in Figure S1A. Note that missingness was heavily concentrated in certain batteries (IADLs and CESD).

**Table 1.**
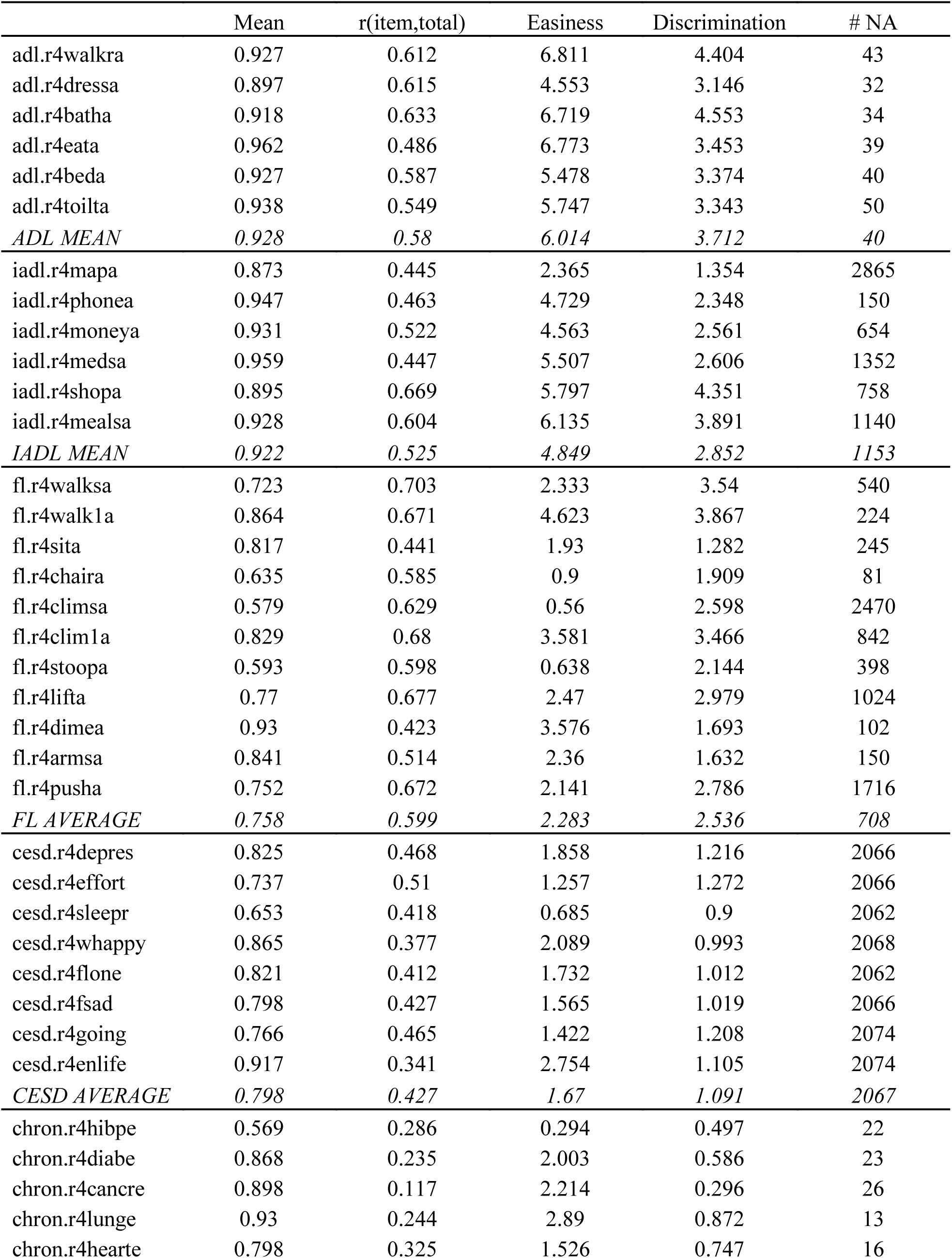
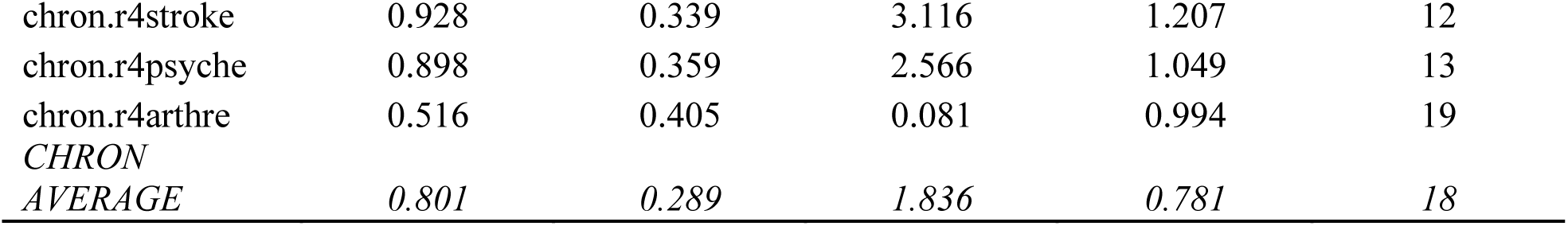
Item statistics for 21384 respondents with health indicators from 1998.

### 2C. Methods

We estimate two parameter logistic (2PL) models. The 2PL model assumes that an individual’s response to an item *i*, *Y*_*i*_, is a function of the individual’s health, *θ*, and two item parameters, *a*_*i*_ and *b*_*i*_:

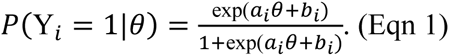

The probability of responding affirmatively (i.e., not having the limitation or health impediment of interest) is determined by the respondent’s level of “health” (represented as *θ*) and the items parameters *b*_*i*_ (the item easiness) and *a*_*i*_ (the discrimination). Note that if the respondent’s latent health (*θ*) was observed, this would reduce to a standard logistic regression problem.^2^ For identification purposes, we assume that health is distributed as Normal[0,1]. Estimates were obtained via the mirt package in R (30). We estimate our models using multiple group maximum-likelihood estimation using an expectation-maximization algorithm and ability estimates expected a-posteriori factor score estimation.

We compare the predictive utility of SRH and BSH using a number of descriptive and non-parametric models. We also consider a series of models related to the prediction of time until death. We consider two baseline models:

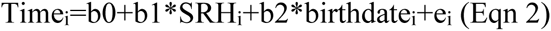

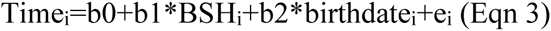

where our general measures of health (BSH in Eqn 2; SRH in Eqn 3) are used to predict time until mortality for those who have recorded deaths as of the wave 11 of HRS data collection. We then consider a model that includes both health predictors:

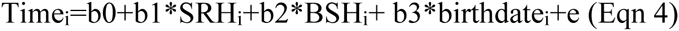

Interest is in the attenuation of the parameters in Eqn4 relative to their baseline values in Eqns 2-3.

## 3. Results

### 3A. The relationship between BSH and the different indicators

Figure 1 focuses on the item characteristic curves. These curves describe the probability of not having some form of ill health (a physical limitation, a chronic condition, or a mental health problem) for a given value of BSH as given by Eqn 1. [Further information on the item parameter estimates is also shown in final two columns of Table 1; individual item characteristic curves are shown in the SI]. We discuss these curves by item battery. ADL items and IADL items tended to be the least prevalent (easiest items). This means that the respondent had to be quite low on the BSH scale before they were unable to perform the ADL tasks. IADL items were fairly similar (i.e., there is substantial within-battery in item functioning for these two batteries).

**Figure 1.**
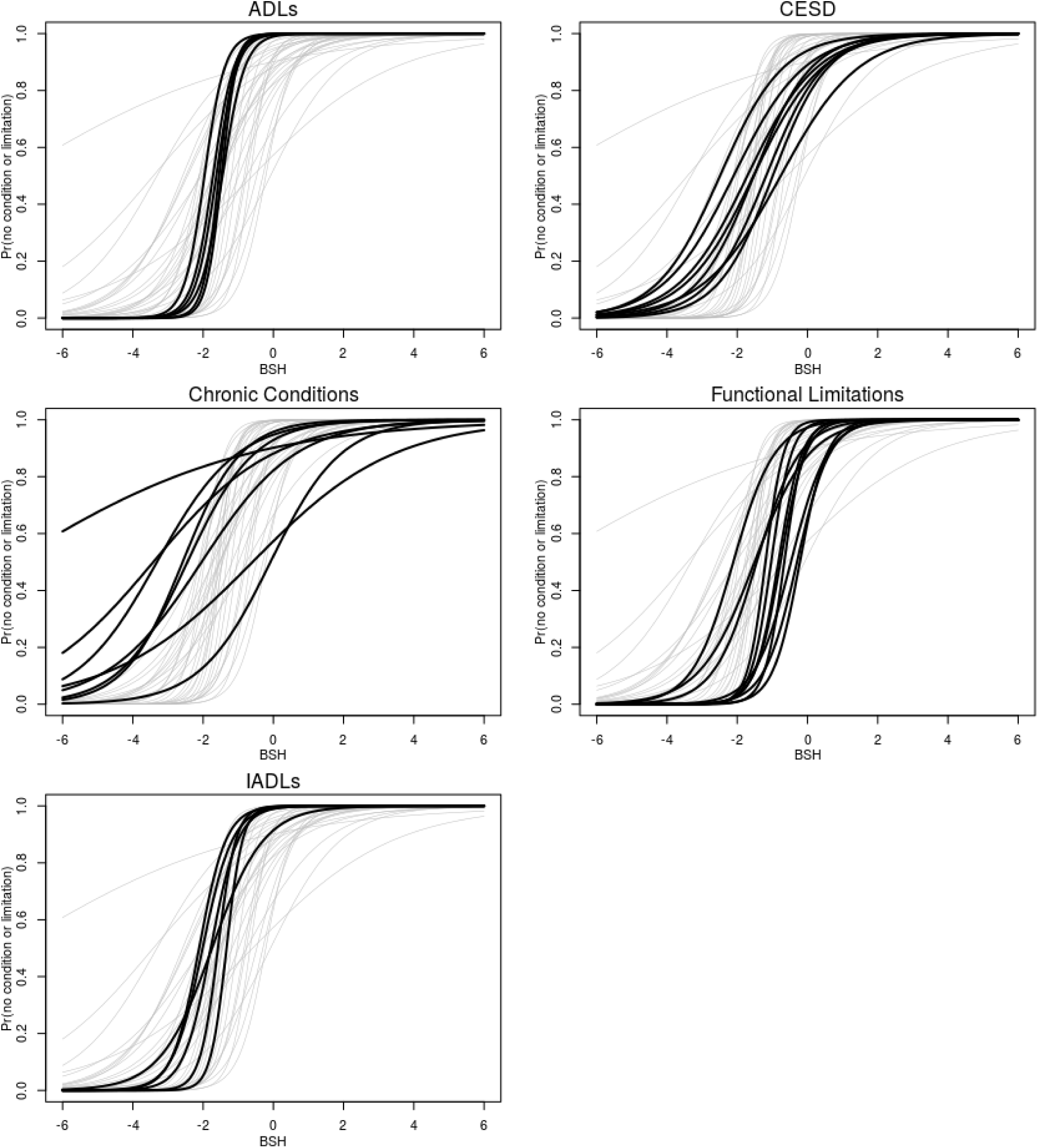
Item characteristic curves for different item batteries.

The functional limitations discriminate over a broad span of health. The items range in easiness from walking one block, on par with the IADL items and something that only those with the least health could not do, to climbing a set of stairs, which was one of the most prevalent indicators of less than optimal health. Taken together, these two items exemplify the broad range of health impediments addressed by this battery. Identifying someone who has a functional limitation as “unhealthy” can mask very large differences between someone who cannot walk up a flight of stairs and someone who cannot walk 1 block.

The CESD items are the least discriminating battery of items, presumably due to the fact that most of the others cover physical functioning and other embodied conditions. Finally, chronic diseases such as cancer and high blood pressure do not do a good job differentiating between the healthy and unhealthy. This is perhaps intuitive, in a sense, since the onset of some disease (e.g., some cancers) are essentially random events (or at least driven by impersonal environmental forces, such as pollutants, rather than human behavior (31)).

Turning to the behavior of the entire set of indicators, we make the following observation. As previously mentioned, there are no items with prevalences less than 0.5. Figure S1B gives an alternative representation. Here, we examine the collective focus of the set of indicators via the test characteristic curve, or the expected number of affirmative items for a given health. We effectively have no indicators that allow us to discriminate amongst the “health” of roughly half of our respondents. Once someone is in the top half of the distribution of health, we are not administering any items inquiring about mild forms of poor health that these respondents are likely to experience. This is a substantial limitation. In the jargon of IRT, this set of indicators has a substantial “ceiling effect” (32).

### 3B. BSH as an improvement on SRH

This ceiling effect is again apparent when we examine individual estimates of health. Figure 2 considers some characteristics of the SRH and (standardized to have M=0, SD=1) BSH measures. Figure 2A shows estimated densities of BSH split by sex in the HRS sample. As in earlier research (33), females have more observable health problems leading to lower BSH. Note that the distributions both show substantial skew due to the “ceiling”. Figure 2B begins to compare the BSH and SRH measures via a boxplot of BSH values for each value of SRH at wave 4. While there is clearly an association between the two measures of health, there is also substantial variation in BSH for each value of SRH. Figure 2C shows the proportion of respondents reporting in each category of SRH (1-excellent through 5-poor) when split by sex. Unlike our BSH measure, there is no sex gradient in the reporting of SRH, a potential indicator of variation in the quality (or at least comparability) of the SRH measure across sex.

**Figure 2.**
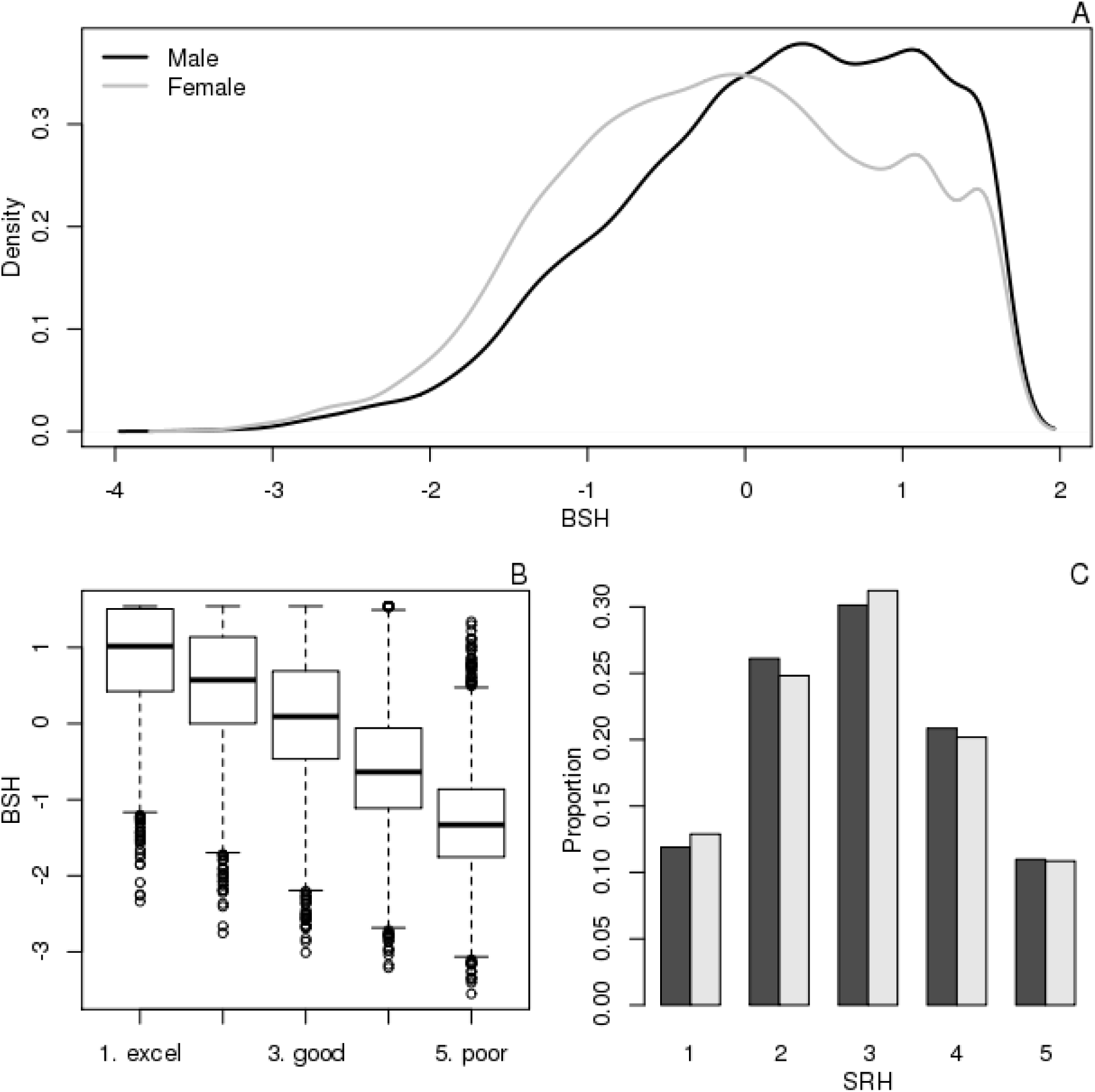
(A) Density of BSH by sex, (B) a comparison to SRH, and (C) proportions of SRH by sex.

As humans have known for millennia, increases in age are associated with declines in health and we now probe the association of our BSH measure with the aging process (Figure 3). Panels 3A and 3B focus on health as a function of age for those who have died as of the 2012 (or Wave 11) of HRS data collection and those who have not. As expected, growing old is associated with declines in the BSH. However, note that the deceased have relatively flat trajectories up until approximately age 70 and then show fairly dramatic decreases. Those who have not died as of the latest wave of data collection tend to be, not surprisingly, much younger, and with better health. Panel 3C focuses explicitly on those who have died (N=8,318) and examines BSH as a function of time before death (defined as time from date of Wave 4 (1998) interview to date of death). As individuals near death, we see steady declines in BSH.

**Figure 3.**
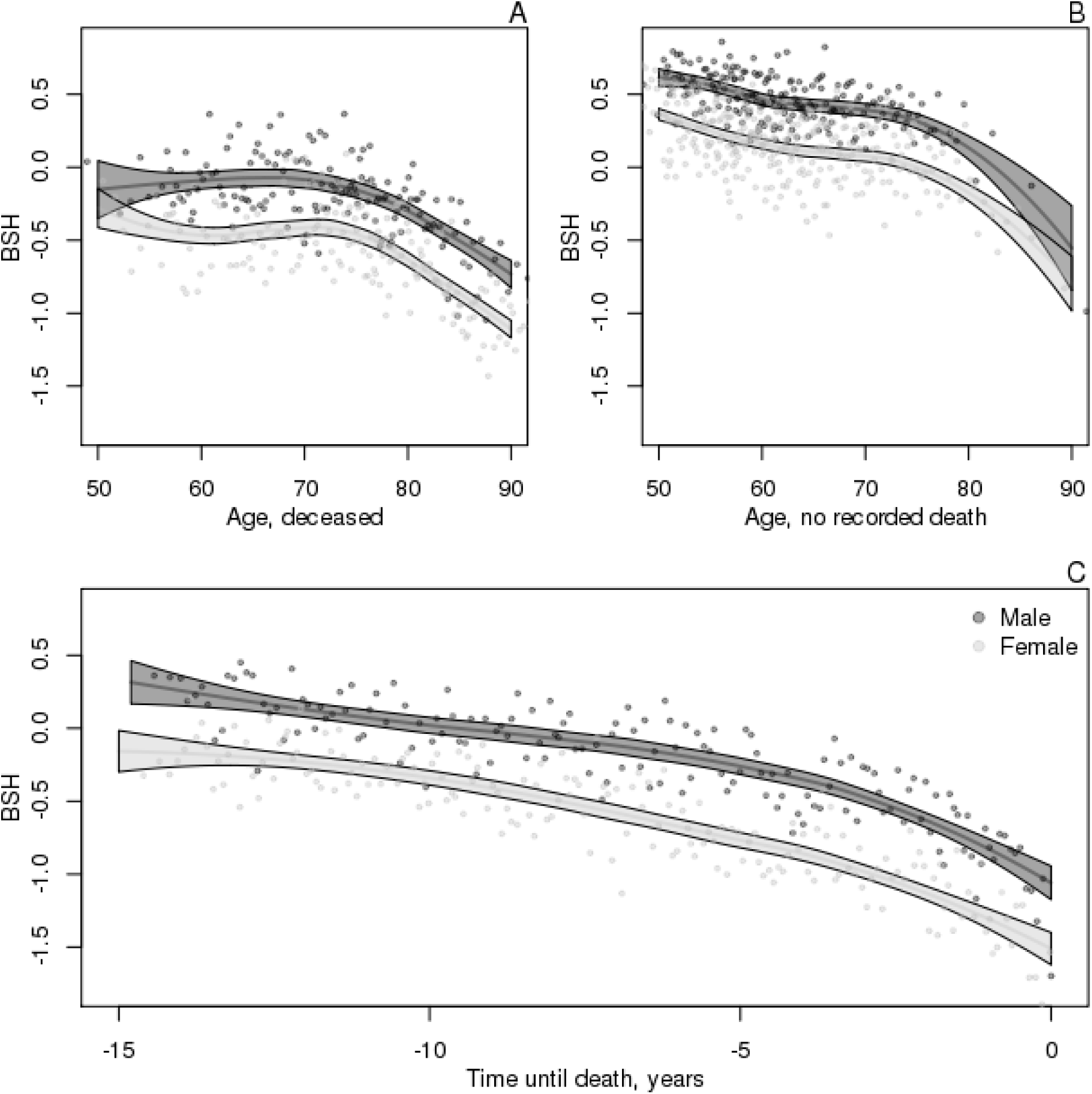
BSH as a function of age (A-for those who die; B-for those with no recorded death) and time before death (C) with binned points each representing approximately 25 people.

We ask now about the association with time until death (the x-axis of Figure 3C) and our two measures of health (BSH and SRH). The distributions of times until death are shown in Figure 4A. Amongst the deceased in our sample, females tend to have slightly less remaining lifespans than do males. We first consider models where BSH *or* SRH is included as a predictor of time until mortality net of date of birth (Eqns 2-3). These are the darker bars in Figure 4B. At baseline, a one SD increase in BSH predicts a 1.2 year increase in time until death. Compared to the base case of poor health, all of the estimates categories of SRH predict increases in time until death of 1.5, 2.5, 3.0, and 3.1 years (respectively for fair, good, very good, and excellent). Table 2 contains r^2^ statistics for these models and suggests that BSH is a slightly superior predictor of time until mortality compared to SRH at baseline.

**Table 2.** Mortality model r^2^ values.

|  | Birthdate only | SRH baseline | BSH baseline | Combined |
| --- | --- | --- | --- | --- |
| All | 0.0613 | 0.1302 | 0.1413 | 0.1547 |
| Males | 0.0513 | 0.1318 | 0.1351 | 0.1537 |
| Females | 0.0767 | 0.1400 | 0.1724 | 0.1789 |

**Figure 4.**
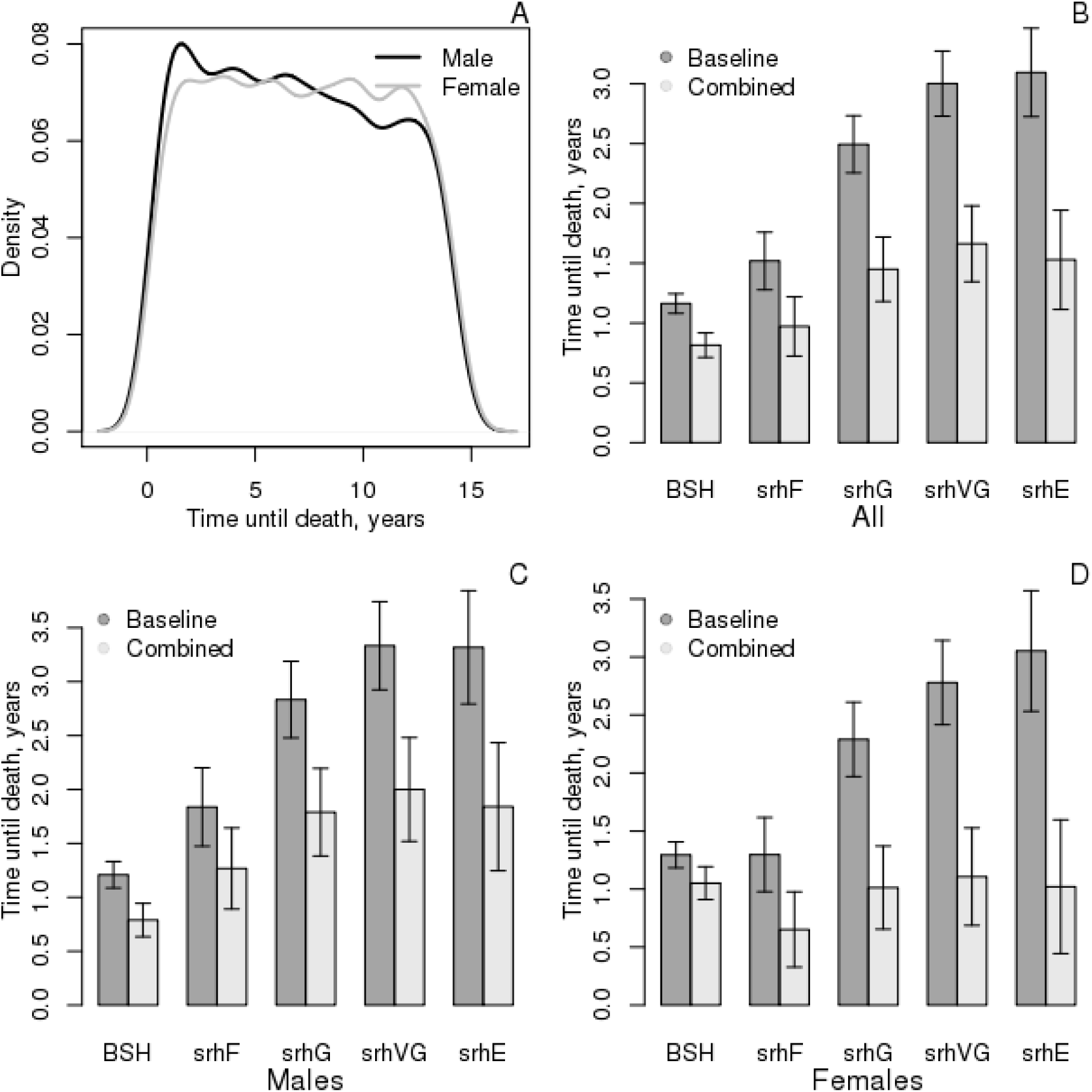
Predictions of time until death using BSH and SRH. (A) Densities of time until death for males and females (N=8318), (B-D) comparison of estimates for all respondents and then split by sex in models of time until death.

We then consider a combined model with both predictors. There is a decrease in BSH, but it is still a strong predictor net of date of birth and self-reported health. Even net of one’s SRH, a one SD increase in the BSH is associated with an additional 8/10 of a year of life. There is, however, a sex gradient to these findings that is explored in panels 4C and 4D. In models without SRH, BSH predicts time until mortality fairly consistently across sex. A 1-SD increase in the measure is associated with an extra 1.2 years of life in males and 1.3 years in females. However, net of SRH, a one SD increase only predicts an extra 0.8 years for males but over a year of extra life for females. Moreover, the effects of the various SRH categories for females are substantially reduced in the combined model. Additionally, the various category effects are similar relative to the effect of poor health (there is little difference between being in fair health versus excellent health). This can be further observed in Table 2 which compares the r^2^ statistics for the models in Figure 4. Note that there is little benefit in terms of r^2^ increase of including SRH for females once BSH is included at baseline.

## 4. Discussion

Using the data from the Health and Retirement Study, we utilize Item Response Theory to combine multiple batteries of health-related indicators (ADLs, IADLs, functional limitations, CESD, and Chronic Conditions) into a single measure of “health” which we describe as broad-sense health (BSH). Looking at the battery-level averages in Table 1, our BSH measure is driven largely by the behavioral modifications (ADLs, IADLs, functional limitations) reported by respondents. Not surprisingly, the depressive symptoms and chronic conditions tend to be both less prevalent and less associated with the overall total. One major finding here is that current questions are quite limited in the sense that they are overwhelmingly measure poor health.

For general populations, researchers may want to consider including items that have higher prevalence (e.g., the existence of back pain; additional items on sleep quality; ability to engage in physical activities that are more strenuous than those in the functional limitations indicators). Developing such indicators is crucial given the simultaneous rise in the number of elderly people who are in relatively good health (34) and the interest in measuring the aging process at in younger populations such as adolescents (35,36). While research focusing on biomarkers will be of keen interest (35,37), survey-based research will continue to be of interest given its cost effectiveness. Although development of such indicators may not be easy, this difficulty is due in part to gaps in current knowledge about the aging process (i.e., what early warning signs at midlife are associated with greater declines in health later in the lifecourse?). Thus, the development and validation of such items may not only offer additional information about the overall health of peoples, but will also potentially help us better understand the nascent health problems which will one day become the types of functional limitations which are the focus of the indicator batteries in HRS.

Despite the limited focus on poor health of the batteries that go into our BSH measure, the proposed measure yielded increased predictive power of time until death relative to SRH. It also reflected a sex-specific story to health declines that earlier research has uncovered. relative improved measurements of global health relative to SRH in some cases. Indeed, we found our measure, Broad Sense Health (BSH) to be a stronger predictor of time until mortality than Self-Rated Health (SRH), especially for females. The results of Figure 3C, suggesting that women are in worse health prior to death (despite the fact that they tend to live longer), has been observed elsewhere (38). The fact that the BSH seems to more reliably uncover sex differences in health and aging vis-à-vis SRH is an important fact.

Research has typically regarded the disablement process as a progression between multiple stages from chronic condition, to symptom, to limitation, to disability and ultimately death (1). Research on the disability has reinforced this viewpoint by showing a near perfect overlap of IADL disability with ADL disability but not the other way around. However, this hierarchy could potentially be scaled via IRT such that we have more information about the “amount” of disablement that must occur to move from an IADL type disability to an ADL type disability. For example, the indicators with the highest prevalence were eating, bathing, and walking across a room (very few HRS respondents could not perform these tasks relative to other indicators of poor health in the HRS). All three of these are ADL items and would, from our theoretical model, imply the most severe disabling limitations. The fourth least easy (most difficult) item in the HRS is ability to prepare a meal, an IADL measurement, which is followed by ability to use a toilet, an ADL measurement. This finding is very surprising and calls into question the precision of ADL and IADL items as currently defined. It is not hard to imagine a world where technological advancement has changed the context of IADL and ADL disability. That is to say the contextual meaning of ADL and IADL measurement were so different in 1998 from the ideal type they were created under that ability to prepare a meal may be more closely reflective of ADL disability than IADL disability. What is certain is that IRT allows us to better understand the severity of different survey disability items. Furthermore, difficulty parameters from functional limitation items reveal a great deal of heterogeneity in what we learn about health from a battery of items that are often combined together. Our results gives estimates of the difference in information about health by items in each battery and suggests that some batteries represent a more unified conceptualization of health while others explore wider ranges of health.

### 4A. Future Research

A crucial methodological limitation is that we are applying unidimensional IRT models to a construct, health, which is presumably multidimensional. Multdimensional IRT models exist (39) and their application may lead to refined BSH measures. Future research examining the utility of multidimensional IRT models would clearly be of interest. Testlet models (40), which allow for dependences between indicators within bundles of items (i.e., “testlets”), are particularly appealing given the fact that the our items are clustered into several well defined batteries (ADLs, chronic conditions, etc). A second area of clear interest would be in cross-cultural studies of health. The MHAS is a parallel study to the HRS focusing on Mexicans. Many of the same batteries are used. The key question in this comparison would be whether the BSH measure is invariant (41) to the cultural context and linguistic changes in the items. Given the sex differences observed here, other measurement variation may well exist. If variation is found, then modifications to more closely align the items, perhaps focusing on their language, may be useful. If no variation is found, then BSH could be used to examine the potential for subjective differences in SRH as a function of cultural context.

### 4B. Conclusion

In this paper we discuss a global health measure based on combining information from many different health batteries. In terms of survey design, this measure suggests that we are generating little information about those individuals who are not in poor health. Given the current state of indicators in HRS, we are not readily able to differentiate between those who are in “fair” and “excellent” health. This is partially a side effect of the focus on the disability process and specific limitations and conditions. Including these in a central manner in studies of disability is rightfully important, but broader measures may be worth considering in an era of increasingly healthy elderly. Despite this shortcoming, we still find our composite indicator to be a superior measure of overall health (e.g., in terms of mortality prediction) than self-reported health. This is especially true for females. Taken together, we think that there is room for improvement in both how we develop indicators asking about health and how we go about operationalizing measures of health once we have asked.

What do we mean by “health”? Those who study disability have a definition of fairly narrow scope. Broader social research largely evades deep thought on the issue by simply asking the respondent to self-report their “health”, whatever that might mean to the respondent. Although the approach we take here was data-driven, we think it illuminates shortcomings in our current conceptions of health (at least to the extent that these conceptions are being operationalized in large-scale surveys). As medical advances allow individuals to live longer and in relatively better health, we think it prudent to begin to expand upon what precisely it is we mean by “health”.

## Acknowledgements

We thank the University of Texas Population Research Center (Grant R24 HD42849) for administrative and computing support; the NICHD Ruth L. Kirschstein National Research Service Award (T32 HD007081-35) for training support; the University of Michigan and Rand Corporation for making the data available to the public, and the members of the Population Health Lab for their helpful suggestions. The contents of this manuscript are solely the responsibility of the authors and do not represent the official views of NICHD, Rand Corporation, Stanford, or the University of Texas at Austin.

## Footnotes

1 We reverse-coded two CESD items (whether the respondent reported being happy or enjoying life).

2 We write Eqn 1 in a slightly non-standard way to emphasize this similarity to logistic regression. Most IRT texts would consider *a*_*i*_ (*θ* + *b*_*i*_).

